# Density-dependent costs and benefits of male mating success in a wild insect

**DOI:** 10.1101/733832

**Authors:** Petri T. Niemelä, Stefano Tiso, Niels J. Dingemanse

## Abstract

1. Social environments are important determinant of fitness, particularly when same-sex local densities shape both mating success and survival costs.
2. We studied how mating success varied across a range of naturally occurring local male densities in wild field cricket males, *Gryllus campestris*, monitored by using fully automated RFID-surveillance system. We predicted that mating success as a function of local density follow a concave pattern predicted by the Allee-effect theory. As increasing density should reduce *per capita* predation and parasitism risk, we predicted that males generally having high mating success in low (versus high) local density live less long. Finally, we predicted that males on average occurred in local densities where their mating success is highest.
3. Male mating success followed a density-dependent pattern predicted by the Allee-effect theory. Males also differed in the local density where their mating success was highest. This variation explained longevity and total fitness: males with high mating success in low local density lived longer and had higher total mating success. Finally, we found no evidence of males occupying local densities in which their mating success is highest.
4. Our study suggest that density-dependent plasticity in mating success is under selection: males having high mating success in low density, but low mating success in high density, lived longer and had higher overall mating success. We thus provide novel insights, with unseen detail, about individual differences in density-dependent mating success and, costs and benefits related to variation in mating success in the wild. Finally, our study also highlights that specific statistical approaches are needed to firmly study the costs and benefits associated with the traits that are repeatedly expressed across range of environments.

## Introduction

Male mating success depends on a diverse array of traits such as colorful or large ornaments (Andersson, 1994), behaviours (Gross, 1996) or acoustic signals (Gerhardt & Huber, 2002). Social environments (e.g., population density) play a major role in shaping the benefits associated with sexually selected traits expressed in males, as they affect a male’s ability to attract females (Bateson & Healy, 2005). For example, the Allee effect posits that average mating success should, at low population densities, increase with increasing population density (Courchamp, Clutton-Brock, & Grenfell, 1999; Stephens, Sutherland, & Freckleton, 1999). Importantly, trade-offs between a male’s investment in reproduction and ability to cope with competition (*sensu* Roff 1992, Dammhahn et al. 2018, Wright et al. 2019) should select for males with relatively high mating success in a high-density environment to have relatively low mating success in a low density environment and *vice versa* (Wright, Bolstad, Araya-Ajoy, & Dingemanse, 2019). Indeed, great tits (*Parus major*) and brown anoles (*Anolis sagrei*), that have high reproductive success (or survival rates) when densities are low have low reproductive success (or survival rates) when densities are high (Calsbeek & Smith, 2007; Nicolaus, Tinbergen, Ubels, Both, & Dingemanse, 2016; Sæther, Visser, Grøtan, & Engen, 2016).

In species where males experience multiple mating opportunities and variation in same-sex density within their life-time, and where the local male density mediates the intensity of competition among males for access to females, the existence of density-dependent mating success may best be studied by adopting a reaction norm approach. This approach enables describing the exact *function* by which mating success varies with density of other males (i.e. intensity of competition) within individuals, and determining how such functions (reaction norms) vary among individual males. Reaction norms are characterized by two main components: First, an intercept represents the average trait value of an individual in a mean-centered environment, and second, a slope represents the change in trait value in unit change on an environmental gradient (Dingemanse, Kazem, Réale, & Wright, 2010; Nussey, Wilson, & Brommer, 2007). Reaction norm slopes can further be linear or nonlinear in nature. The components of reaction norms can be under selection independently from each other (Nussey et al., 2007) and, if they are correlated, the selection pressures acting on one component (e.g. slope) might be affect the selection gradient estimates of other components (e.g. intercept) if not taken into account. If males vary in their density-dependent payoffs, we expect reaction norm slopes, which describe how mating success changes as a function of local density *within* an individual male, to cross, such that some males have highest mating success in high-density environments and others instead have highest mating success in low density-environments. Importantly, traits facilitating mating success are often costly because they are either energetically expensive to produce (Scharf, Peter, & Martin, 2013) or increase the risk of predation or parasitism (Godin & McDonough, 2003; Scharf et al., 2013; Zuk & Kolluru, 1998). Additionally, these costs might vary as a function of population density and individual-specific traits (Cote, Dreiss, & Clobert, 2008; Knell, 2009; Moorcroft, Albon, Pemberton, Stevenson, & Clutton-Brock, 1996), thereby leading to selection favoring males that are successful in a specific local density (e.g. high or low local density). Quantifying individual-level reaction norms would thus allow us to study survival selection (i.e. costs) and fitness selection (i.e. benefit) acting on the shape of the reaction norm describing the change in mating success across a gradient of local male density. Furthermore, the presence of individual-level density-dependent costs and benefits related to mating success is predicted to also lead to adaptive phenotype-environment matching, where individuals seek out densities where they have the highest mating success (Edelaar, Siepielski, & Clobert, 2008).

Ideal model species to study density-dependent variation in mating success, and the costs and benefits associated with this variation, are those forming dynamic social aggregations, such as field crickets (*Gryllidae*). Field crickets, *Gryllus campestris*, live close to each other in small “territories” (a small open area in grassland vegetation), with a burrow, from which males signal acoustically to attract mating partners (Rodríguez-Muñoz, Bretman, Slate, Walling, & Tregenza, 2010; Rodríguez-Muñoz, Bretman, & Tregenza, 2011). Males, moreover, inform their presence to other males via acoustic signaling and these signals also mediate competition between males (Gerhardt & Huber, 2002). Males also frequently change burrows (Niemelä & Dingemanse, 2017; Niemelä, Lattenkamp, & Dingemanse, 2015; Rodríguez-Muñoz et al., 2011), which makes male densities dynamic but also potentially enables adaptive phenotype-environment matching (see above). Thus, the ecology of our model species facilitates the study of density-dependent male mating success using a reaction norm framework detailed above.

Here we study, first, whether local male density affects male mating success at the population level. We then estimate whether the effects of local density on mating success vary among-individuals, i.e. whether there is individual variation in mating success reaction norms as a function of density. Finally, we study whether variation in density-dependent mating success explains variation in longevity and total reproductive success and, whether males select environments where their expected mating success is highest (adaptive phenotype-environment matching). As a model, we use wild field crickets of the species *G. campestris*; this species group is traditionally used to study sexual selection (Bailey & Zuk, 2009; Cade, 1981; Cade & Cade, 1992; Hunt et al., 2004; Rodríguez-Muñoz et al., 2010; Tregenza, Simmons, Wedell, & Zuk, 2006; Tregenza & Wedell, 2002). As a proxy for mating success, we used hourly mating success, defined as the hourly probability that a focal male is detected with a female in its burrow. This probability positively predicts mating success in our model species (Rodríguez-Muñoz et al., 2010, 2011). We predicted a concave shape of the reaction norm (describing hourly mating success as a function of the local density) at the population level (Courchamp et al., 1999; Stephens et al., 1999). Since in our model species, high quality males have been shown to pay mortality costs, potentially because the presence of a female at his burrow increase male mortality through predation or because intrinsic costs related to intense signaling (Hunt et al., 2004; Rodríguez-Muñoz et al., 2011), we predicted that individuals with higher average daily mating success lived shorter. Since *per capita* risk of predation should be lower when local densities are high (Kacelnik, Krebs, & Bernstein, 1992; Ryan, Tuttle, & Taft, 1981), we further predicted that males exhibiting high mating success in high density environments lived longer. Finally, we predicted phenotype-environment matching to exist (Edelaar et al., 2008).

## Methods

The data was collected in a 50mx50m field plot near the Max Planck Institute for Ornithology (Seewiesen, southern Germany: 47°58□35.5□N, 11°14□04.5□E), from 8 May to 6 June 2015. The study area was delimited by transparent plexiglass panels, high enough to avoid crickets to enter or exit the area (Niemelä & Dingemanse, 2017). Our enclosure nevertheless allowed ample movement because of its large relative size: common field crickets typically move only over short distances (on average 35 m) following post-wintering hibernation (Ritz & Köhler, 2007). In early spring, all subjects emerged as adults within our study area from natural (selfmade) burrows, where they had overwintered, and from which they could move freely. In this species, nymphs become active in early spring (March) and stay in close proximity of their burrow (c. 10-20 cm distance). After transforming into mature adults (the stage at which our data was collected), both sexes start searching for mating partners (Niemelä et al., 2015; Rodríguez-Muñoz et al., 2011). Given that crickets spend the majority of their time in, or nearby, burrows, they can easily be trapped, marked, and their behavior recorded for their entire adult lifespan in great detail (Fisher, David, Tregenza, & Rodríguez-Muñoz, 2015; Niemelä & Dingemanse, 2017; Niemelä et al., 2015; Rodríguez-Muñoz et al., 2010, 2011).

### Data collection

To mark individuals, they were trapped using custom-made traps consisting of two parts: a 30-cm-long tube that was inserted into the entrance of the underground part of the burrow, and a cylinder-shaped chamber, in which the cricket would fall after climbing the tube while attempting to leave the burrow (Niemelä & Dingemanse, 2017). Each field cricket was marked after reaching sexual maturation (early May) (n=90; 59 females, 31 males) with a small circular numbered “bee” tag (diameter 2.5 mm), and fitted with a RFID-tag (Passive Integrated Transponder: length 8 mm, diameter 1.4 mm; Trovan Ltd. Isle of Man). Tags were attached to the pronotum using cyanoacrylate adhesive, which has been also used in other studies of our model species to attach objects of similar size and weight (Fisher et al., 2015; Rodríguez-Muñoz et al., 2010, 2011). A preliminary experiment from 2014 showed that that RFID-tags do not affect survival in the wild over the period of 20 days for tagged versus untagged individuals (n=20 individuals in each treatment): (daily survival probability (95% Credible Intervals) = 0.99 (0.96-1.00) versus 0.98 (0.95-1.00), respectively; Niemelä et al. Unpublished data).

We placed one circular RFID-antenna (Trovan Ltd. Isle of Man, diameter: 40 mm) at every burrow in the study area few days before the crickets were predicted to transform into an adult. The total number of monitored burrows was 124 throughout the season. The total number of antennas was 94, the difference between number of burrows and number of antennas resulting from crickets creating new burrows when old ones were abandoned or destroyed. In such cases, we moved the antennas from the abandoned/destroyed burrows to the new one, causing the number of burrows throughout the season to be higher than the number of antennas. The antennas were placed so that they covered the entrance of the burrows: to enter or exit the burrow, crickets had to walk through the antenna. The antennas read the identity of the RFID-tags with a date/time (reading resolution of 1 second) stamp each time a PIT-tagged cricket entered reading range (~1-2 cm beyond both lateral sides of the antenna). Data was sent wirelessly, through main reader units (each controlling for 20-25 antennas) to a computer in nearby building (~500 meters away). The system thus collected spatiotemporal data on the location of each individual in the study area.

Preliminary tests, made by comparing 21, 30-minute long, videos with RFID readings (recorded from 15 different burrows) showed that the RFID-antennas missed a reading in only 6.5% of cases where the cricket was within the antenna range (Niemelä et al. Unpublished data). These missed data should not cause major bias given that each cricket was read hundreds of times per day (with 1 second interval) when occupying a burrow, implying that presence at a burrow would typically be detected by the system. The total number of readings by the RFID-system was 1,846,831.

### Estimating local density and mating success

For hourly mating success, for each hour (*t*), we scored whether the focal male was detected with a female in its burrow or not (1/0; binary data). For each burrow and hour, we calculated local male density defined as the number of males/m^2^ (which included the focal male) during the previous hour (*t*-1). We focused on local male density since males compete with each other, mainly acoustically, for mating opportunities (Gerhardt & Huber, 2002). Females are not acoustically active and thus, female density may be hard to estimate by crickets of either sex, and responses to it hard to evolve, and was therefore not analyzed. For local male density, we chose 500 centimeters as our upper spatial range limit since male crickets generally stop responding to the acoustic signals of others way before that limit (Cade 1981, Simmons 1988, Hissman 1991). Since we did not have strong prior information about the exact distance where such social environment effects might be of importance, we calculated density using different radius criteria (100, 200, 300, 400 & 500 centimeters), and asked which one best explained the data. Then we selected the best-fitting distance (see “Statistical methods”) for analyses presented in the main text. Notably, there were 1, 7, 18, 18 & 18 focal burrows with 100, 200, 300, 400 & 500 centimeter radius, respectively, that had surface areas partly outside (i.e., overlapping with the border of) our study plot. For such burrows-radius combinations, we performed our density calculation using only the surface area inside our study plot. We also statistically estimated whether an “edge” effect was present by including distance between the edge and the focal burrow as a covariate in our models; edge effect were not detected (supplementary table 1), and therefore not included in the models presented in the main text.

We only used data from individuals with known age, i.e. males that were marked directly following maturation. This practically meant that we removed males from the focal male data set that were found, and marked, in the middle/end of the season (specifically, days 15-27 after the maturation of the first individual; n = 7 males with unknown age). Removal of males with unknown exact adult age was important as age affects many traits in our model species, including the sexual signaling and attractiveness (Rodríguez-Muñoz et al., 2019; Verburgt, Ferreira, & Ferguson, 2011), and thus had to be controlled for in our models. Notably, those males with unknown exact adult age were included when calculating local density. Our final sample size was 7603 (hourly) data points for 24 males over a 30-day period, occupying 76 different burrows fitted with readers. Each male visited on average 5.75 (SD: 2.74) unique burrows and made 10.03 (SD: 7.29) burrow changes. In other German field cricket populations, ~70% of the adult males live less than 20 days (Ritz and Kohler 2007). This means that our sampling period of 30 days (36 days for longevity data) covered the majority of the lifespan of our model species.

## Statistical methods

### Model selection and population level mating success

As a first step, we ran five separate univariate mixed effects models for the mating success, one for each defined radius (100, 200, 300, 400 & 500 centimeters) considered (see above). In all models, we included age, squared age, among-subject centered male density, squared among-subject centered male density within-subject centered male density and squared among-subject centered male density as continuous covariates. Age was mean-centered and variance-standardized. Local density was mean-centered and variance-standardized before among- and within-subject centering was applied. Covariates were standardized to facilitate model convergence. Among- and within-subject centering of the local density allowed us to separate within-individual variation in mating success as a function of local density (i.e., within-individual plasticity) from patterns caused by individual differences in average density experienced (an among-individual level phenomenon) (van de Pol & Wright, 2009). We also included individual identity, date identity and burrow identity as random effects to estimate variation in response variable caused by spatiotemporal patterns and individual. Model selection, where the models with different radius from the focal burrow were being compared, was based on AIC-values so that ΔAIC>7 was considered as a significant difference in fit (Burnham, Anderson, & Huyvaert, 2011). All models had identical fixed and random effects structure.

### Individual variation in mating strategies as a function of local density

As a second step, we ran one bivariate random slopes mixed effects model, where we included longevity as the first and hourly mating success as a second response variable. We did this to estimate individual-level BLUPs (i.e. best linear unbiased predictors; (Henderson, 1975)) for longevity and for the average hourly mating success (i.e. intercept) and linear and non-linear density-dependent plasticity in hourly mating success to be used to estimate survival and fitness selection gradients (see below). This random slopes model allowed us to estimate whether individuals differed in i) average hourly mating success, ii) linear and non-linear density-dependent plasticity in hourly mating success and, iii) whether the average hourly mating success and density-dependent plasticity in mating success covaried with each other and with longevity (Nussey et al. 2007, Dingemanse et al. 2010). To estimate abovementioned components of the reaction norm, we included the interaction between the within-subject centered local male density (linear and quadratic terms) and individual identity in the model as a random term. Otherwise, we used the same fixed and random effects structure as in the first step (see above). Thus, the reaction norms and their associated BLUPs are controlled for spatiotemporal variation, age effects and the variation in average local density among-individuals.

### Costs and benefits related to mating success

As a third step, we modeled survival selection gradients for the male’s average hourly mating success (i.e. intercept) and for the linear and quadratic density-dependent plasticity in hourly mating success (i.e. slopes of the reaction norm) by using general linear model. For the model we included relative longevity (mean-standardized days survived) as response variable and standardized individual level BLUPs for the intercept and the linear and quadratic slopes of the reaction norm describing the hourly mating success as a function of local density as covariates. This way any confounding factors, e.g. correlations between intercept and slopes of the reaction norm, are controlled for.

We also estimated selection gradients as described above for total fitness, defined as total mating success throughout the study period (i.e. the life-time of the individual). Total mating success was estimated by estimating the total amount of hours alive for each male and then multiplying this by the hourly mating probability of each male. This type of fitness proxy correlates with total reproductive success in our model species in the wild (Rodríguez-Muñoz et al., 2011).

### Phenotype-environment matching

Finally, we estimated whether individual variation in average hourly mating success (i.e. intercept) and linear and quadratic density-dependent plasticity in hourly mating success explained individual variation in experienced density. We did so using a general linear model which included each individual’s average local density (BLUP) as the response variable and each individual’s BLUP for the intercept (average hourly mating success) and the linear and quadratic slopes (i.e. plasticity in hourly mating success) of the reaction norm as covariates. The BLUPs for local density were estimated by using mixed effects model with mean-centered local density as a response variable and male identity, burrow identity and date identity as random effects.

The usage of BLUPs has been criticized in the literature because with low number of repeats (i.e. 2-3 measurements/individual), BLUPs are estimated with large error (Houslay & Wilson, 2017). However, the BLUPs for reaction norm components and for local density in our study are estimated with the average sample size of 316.8 (SD; 124.36) measurements (i.e. hours) for each individual, effectively making such problems obsolete in our case.

All models were run in the R statistical environment (R core team, 2014). For model selection and population level effects, selection gradients and phenotype environment matching, we used the package lme4 (Bates, Maechler, Bolker, & Walker, 2015) with binomial and Gaussian link functions. To estimate individual level reaction norms for hourly mating success across local density and correlations between reaction norm components and longevity, we used the MCMCglmm package (Hadfield, 2010) with a Gaussian (for longevity) and binomial (for hourly mating success) link functions. For the MCMCglmm model, we used inverse gamma-distributed priors; the model was run with 1,530,000 iterations with 30,000 burn-in and sampling rate of 1000, resulting in low autocorrelation between samples.

## Results

### Model selection and population level hourly mating success

Of the models fitting density using 100-500 centimeter radiuses, the one where local density was estimated using a 300-cm radius from the focal individual’s burrow fitted the data best (Table 1). For further analyses, this 300-cm radius was used to define local density. This best fitting model showed a significant positive linear effect of age on hourly mating success (Table 1); the population-average hourly mating success increased with age. Moreover, there was a negative linear and non-linear effect of local density on hourly mating success (Table 1), which followed the shape predicted by an Allee-effect (Courchamp et al., 1999; Stephens et al., 1999). That is, population-average hourly mating success increasing from low to intermediate local densities and then declined with further increases in density (Figure 1).

**Table 1.** Sources of variation in hourly mating success for different local densities (i.e. 100-500 centimeter radius from a focal burrow) and, without local density; we present fixed (β) and random (σ^2^) parameters with 95% Credible Intervals, as well as model AIC-values, derived from univariate mixed effect models. Estimates and 95% Credible Intervals are derived by simulating the model 1000 times.

|  | Density 100 | Density 200 | Density 300 | Density 400 | Density 500 | No Density |
| --- | --- | --- | --- | --- | --- | --- |
| Fixed effects | $\beta$ (95% CI) | $\beta$ (95% CI) | $\beta$ (95% CI) | $\beta$ (95% CI) | $\beta$ (95% CI) | $\beta$ (95% CI) |
| Intercept | -3.95 (-5.84, -2.11) | -3.46 (-5.17, -1.69) | -2.90 (-4.74, -1.13) | -2.83 (-4.70, -0.87) | -2.71 (-4.55, -0.87) | -3.23 (-5.10, -1.63) |
| Age | 2.28 (1.35, 3.28) | 2.43 (1.47, 3.48) | 2.30 (1.24, 3.42) | 2.06 (1.15, 3.07) | 2.19 (1.17, 3.23) | 2.13 (1.14, 3.11) |
| Age<sup>2</sup> | -0.01 (-0.29, 0.29) | <-0.01 (-0.27, 0.27) | -0.04 (-0.30, 0.24) | -0.05 (-0.33, 0.23) | -0.04 (-0.31, 0.23) | -0.04 (-0.29, 0.24) |
| Density_among | -3.35 (-6.29, -0.52) | 2.39 (0.06, 4.61) | 2.16 (0.26, 4.11) | -0.09 (-2.00, 1.78) | 0.37 (-1.44, 2.06) | - |
| Density_among<sup>2</sup> | 4.91 (1.55, 8.20) | 2.42 (-1.60, 6.61) | -0.70 (-3.50, 2.18) | -2.15 (-5.47, 0.80) | -1.60 (-3.83, 0.56) | - |
| Density_within | 0.08 (-0.08, 0.21) | 0.03 (-0.11, 0.18) | -0.33 (-0.46, -0.18) | -0.36 (-0.50, -0.20) | -0.04 (-0.23, 0.15) | - |
| Density_within<sup>2</sup> | -0.13 (-0.19, -0.08) | -0.12 (-0.20, -0.04) | -0.25 (-0.33, -0.16) | 0.06 (-0.02, 0.13) | -0.25 (-0.35, -0.15) | - |
| **Random Effects** |  |  |  |  |  |  |
| Variance | $\sigma^2$ (9% CI) | $\sigma^2$ (SD) | $\sigma^2$ (SD) | $\sigma^2$ (SD) | $\sigma^2$ (SD) | $\sigma^2$ (SD) |
| Residual | 3.29 (fixed) | 3.29 (fixed) | 3.29 (fixed) | 3.29 (fixed) | 3.29 (fixed) | 3.29 (fixed) |
| Burrow | 15.78 (12.21, 20.55) | 15.62 (12.29, 19.79) | 15.78 (12.24, 20.09) | 16.13 (12.32, 20.34) | 15.20 (11.20, 19.10) | 15.44 (12.06, 19.42) |
| Date | 9.14 (7.02, 12.76) | 9.25 (7.21, 13.05) | 9.55 (7.42, 12.97) | 8.91 (6.90, 12.33) | 8.87 (6.94, 12.00) | 8.52 (6.67, 11.57) |
| Individual | 1.84 (1.36, 2.67) | 1.40 (1.14, 1.86) | 1.75 (1.34, 2.45) | 2.55 (1.98, 3.44) | 2.68 (2.10, 3.66) | 2.87 (2.26, 3.77) |
| **Model fit** |  |  |  |  |  |  |
|  | AIC | AIC | AIC | AIC | AIC | AIC |
|  | 4446.2 | 4457.8 | **4404.2** | 4456.1 | 4445.3 | 4470.4 |

**Figure 1.**
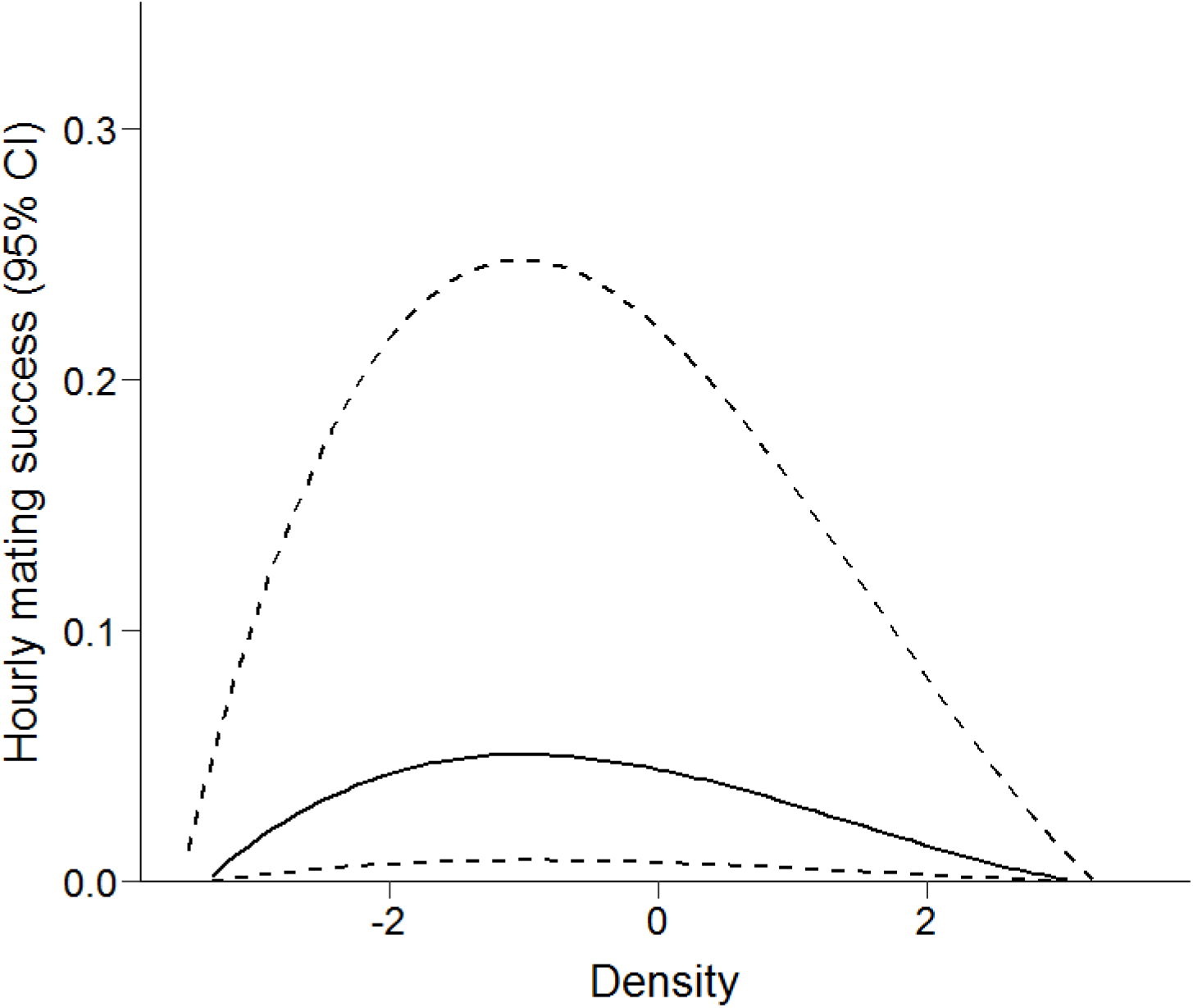
Population level mode for hourly mating success across mean centered local densities within 300 centimeter radius from the focal burrow (black solid line). Black dashed lines refer to 95% Credible Intervals. The regression line and Credible Intervals are constructed of data-scale predictions generated by simulating the univariate glmer-model 1000 times.

### Individual variation in mating success as a function of local density

There was significant individual variation in average hourly mating success (reaction norm intercept; σ^2^_i_ (95% CI): 4.65 (1.91-8.26) and both linear (4.66 (1.56-9.18)) and non-linear (1.34 (0.43-2.59) (Figure 2) density-dependent plasticity in hourly mating success (i.e. linear and nonlinear slopes, respectively).

**Figure 2.**
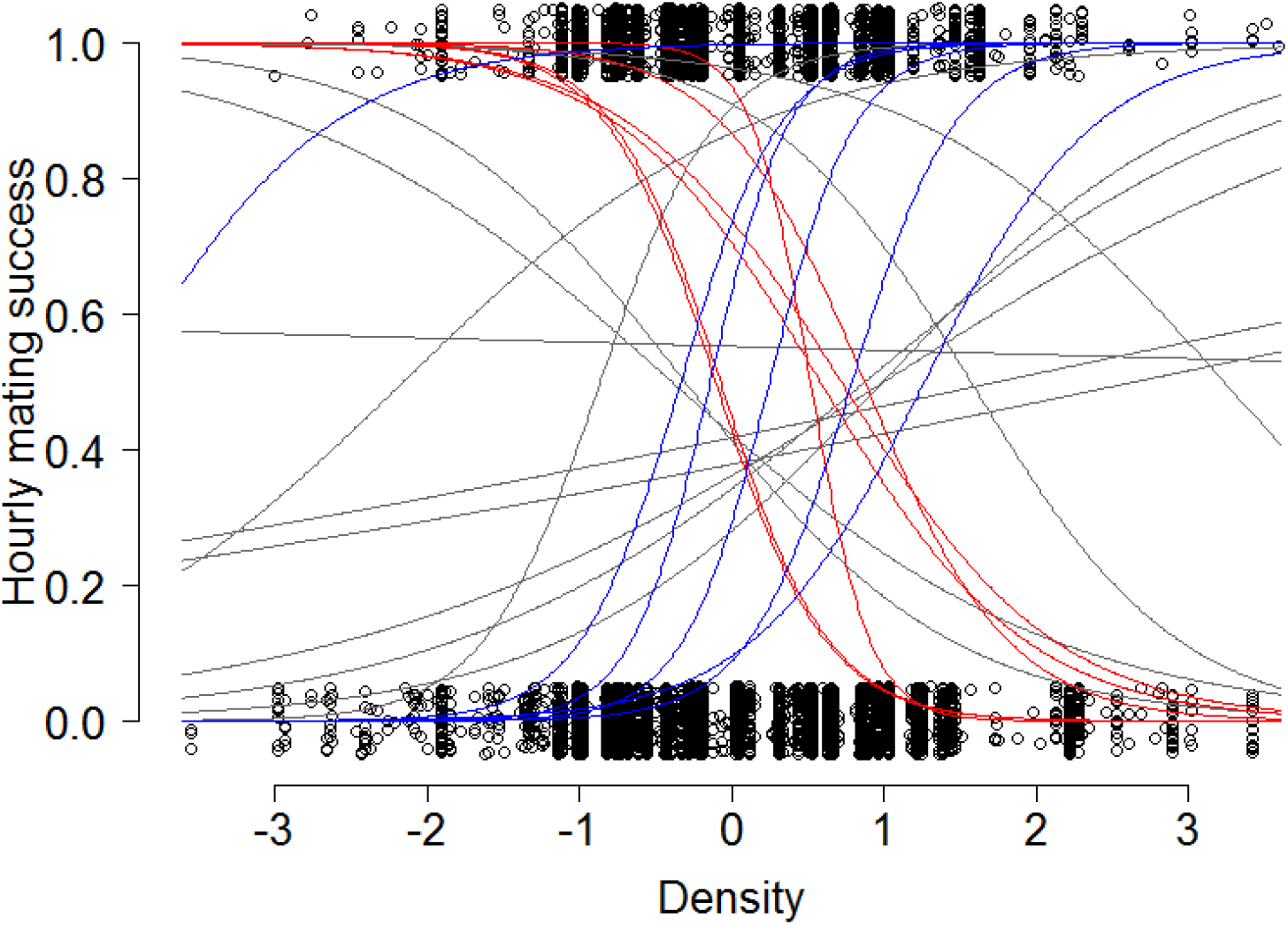
Predictions of individual level reaction norms for hourly mating success as a function of mean-centered density. Red lines refer to 6 individuals with steepest negative linear slopes, blue lines refer to 6 individuals with steepest positive linear slopes and the 12 grey lines refer to rest of the individuals. The reaction norms are constructed from the BLUPs generated by the random slopes MCMCglmm-model. Open circles represent the raw data.

### Costs and benefits related to mating success

Longevity was negatively correlated with an individual’s linear density-dependent plasticity in hourly mating success (*r*_i_ (95% CI): −0.68 (−0.86; −0.09)) (Figure 3). Male’s average hourly mating success (i.e. intercept) and non-linear density-dependent plasticity in hourly mating success were not correlated with longevity (*r*_i_(95% CI): 0.22 (−0.22; 0.66) & −0.23 (−0.73; 0.34), respectively). Those findings implied that individuals with high hourly mating success in low density, but low hourly mating success in high density, live longer (Figure 2, Figure3). This was also confirmed by the survival selection gradient model showing that only the individual’s linear change (linear slope) in hourly mating success across density gradient was under negative selection (Table 2a, Figure 3). Average hourly mating success (intercept), linear and non-linear density-dependent plasticity in hourly mating success were not correlated (*r*_i_(95% CI): intercept & linear slope; −0.04 (−0.59; 0.42), intercept & non-linear slope; −0.35 (−0.68; 0.47), linear & nonlinear slope; 0.60 (−0.01; 0.87)). This indicates that males doing well under low density (i.e. negative linear slopes) do not have higher average hourly mating success compared to males that do well under high density (i.e. positive linear slopes).

**Figure 3.**
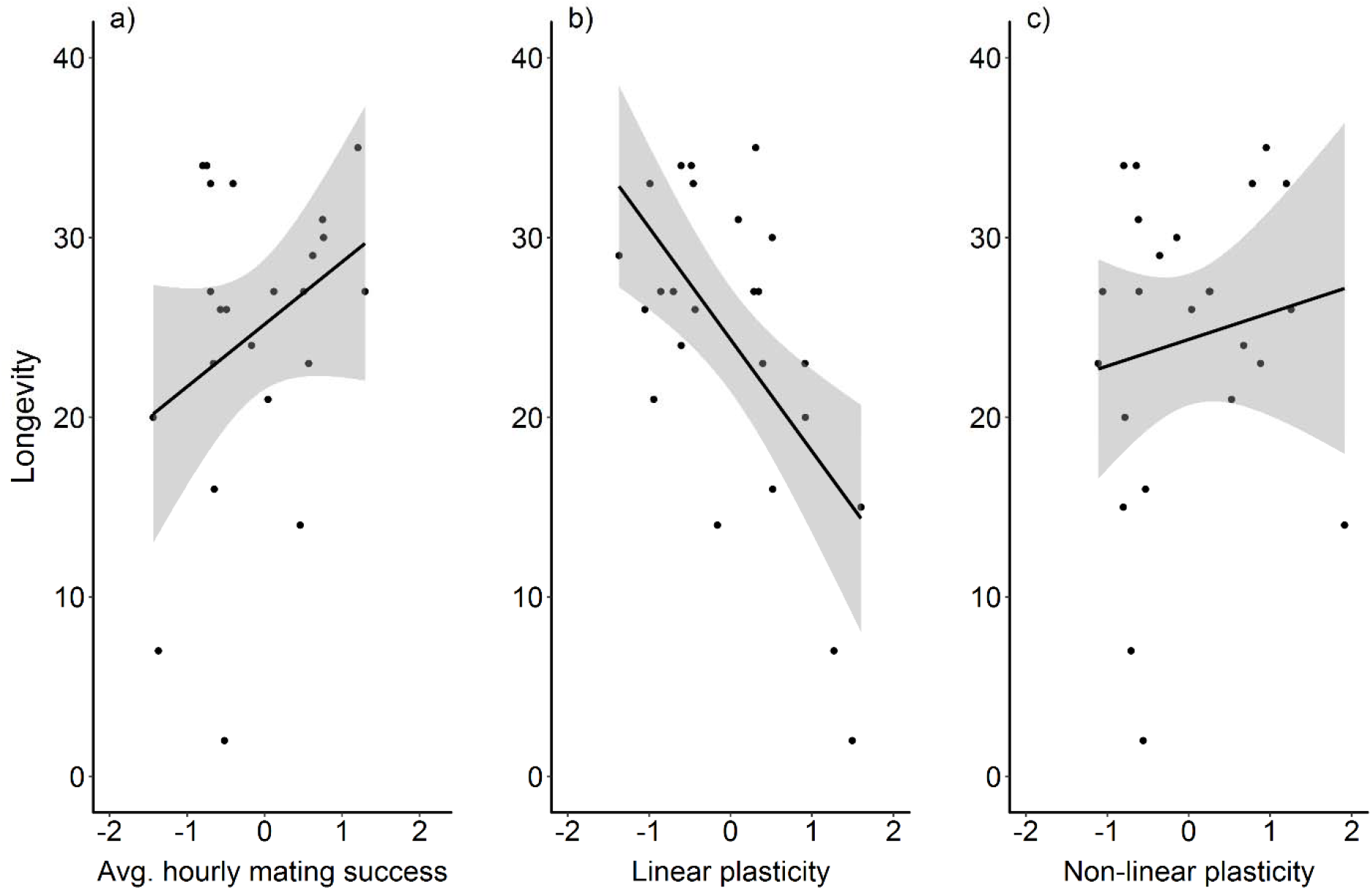
Longevity plotted as a function of individual level estimate (BLUP) for a) average hourly mating success and b) linea r and c) non-linear density-dependent plasticity in hourly mating success. Shaded grey area refers to standard error. The black dots are residuals from the models where each focal component of the reaction norm (BLUP) is set separately as a response variable (i.e. 3 models) and the other two components are set as covariates. Thus, residuals are fully independent components of each BLUP, i.e. any non-independence between reaction norm components have been controlled for. The black solid line is a regression line describing the association between y and x.

**Table 2.** Sources of variation in a) longevity and b) total mating success explained by standardized average hourly mating success and linear and non-linear density-dependent plasticity in hourly mating probability. We present fixed (β) parameters with standard errors, as well as test statistics (*T*-value) with accompanying *P*-values, derived from a general linear models. Since we use relative longevity and total mating success, and standardized covariates, the β-estimates represent survival and fitness selection gradients, respectively.

| a) | $\beta$ (SE) | $T$-value | P-value |
| --- | --- | --- | --- |
| Longevity intercept | 1.00 (0.057) | 17.407 | <0.001 |
| Hourly mating success | 0.03 (0.060) | 0.538 | 0.597 |
| Linear plasticity | -0.26 (0.069) | -3.694 | 0.001 |
| Non-linear plasticity | 0.06 (0.069) | 0.872 | 0.394 |

| b) | $\beta$ (SE) | $T$-value | P-value |
| --- | --- | --- | --- |
| Total mating success intercept | 1.00 (0.073) | 13.792 | <0.001 |
| Hourly mating success | 0.48 (0.076) | 6.330 | <0.001 |
| Linear plasticity | -0.20 (0.087) | -2.315 | 0.031 |
| Non-linear plasticity | 0.02 (0.087) | 0.178 | 0.860 |

Total mating success was positively associated (i.e. positive selection gradient) with the male’s average hourly mating success and negatively associated (i.e. negative selection gradient) with an individual’s linear density-dependent plasticity in hourly mating success (Table 2b, Figure 4). This indicates that males that have high hourly mating success, have also high total mating success. Moreover, males that are successful in attracting mating partners at low density environments had a higher total mating success compared to males that are successful in high density environments.

**Figure 4.**
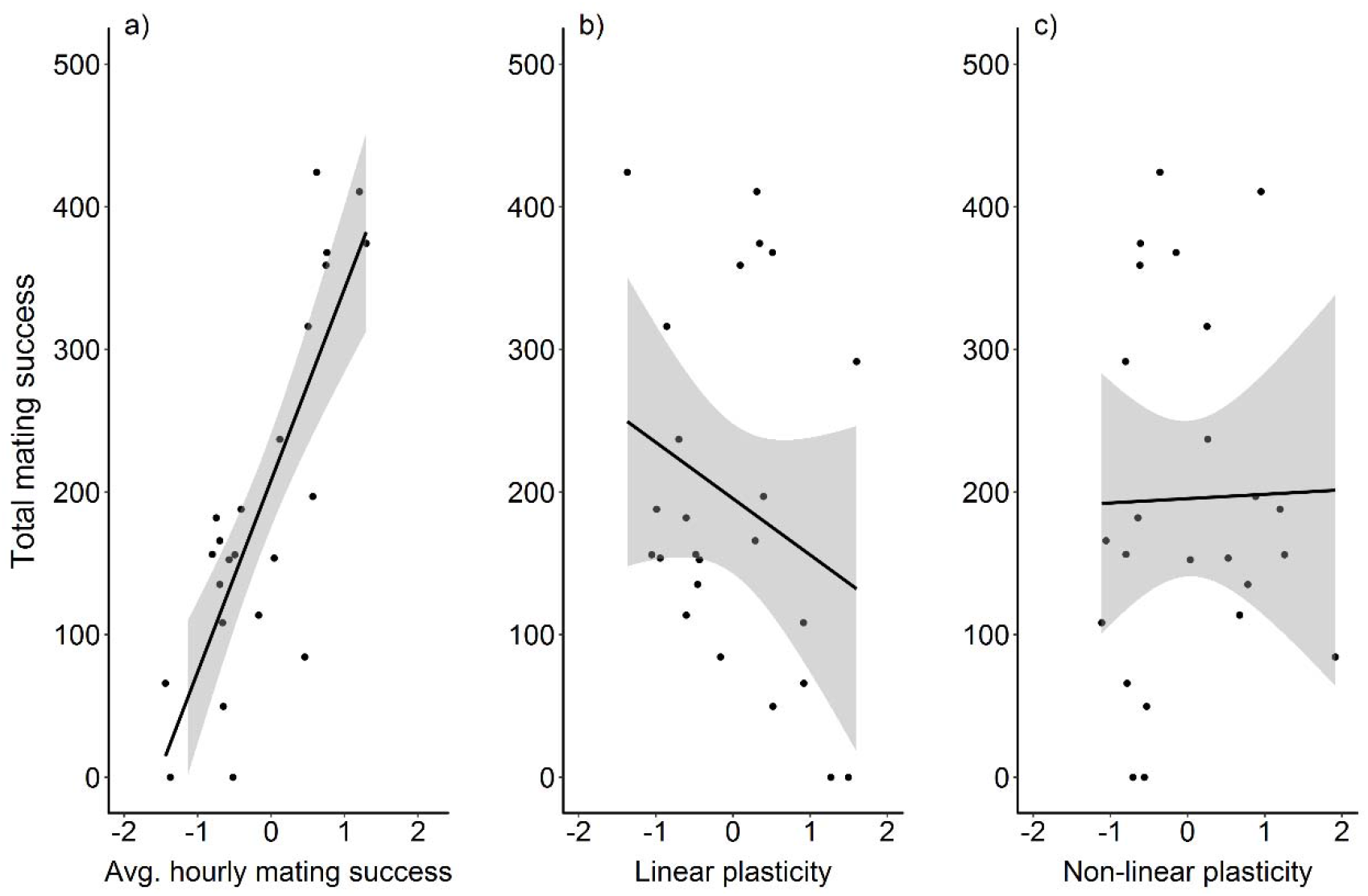
Total mating success plotted as a function of individual level estimate (BLUP) for a) average hourly mating success and b) linear and c) non-linear density-dependent plasticity in hourly mating success. Shaded grey area refers to standard error. The black dots are residuals from the models where each focal component of the reaction norm (BLUP) is set separately as a response varia ble (i.e. 3 models) and the other two components are set as covariates. Thus, residuals are fully independent components of each BLUP, i.e. any non-independence between reaction norm components have been controlled for. The black solid line is a regression line describing the association between y and x.

### Phenotype-environment matching

Males were repeatable in local density (R (95% CI): 0.29 (0.17-0.43)). However, neither average hourly mating success (intercept) nor level of density-dependent plasticity (whether linear or nonlinear) predicted an individual’s average local density (Table 3). This means that males did not seem to select local density environments according to where their expected mating success would be highest.

**Table 3.** Sources of variation in individual level average density (BLUP) explained by standardized average hourly mating success and linear and non-linear density-dependent plasticity in hourly mating success. We present fixed (β) parameters with standard errors, as well as test statistics (*T*-value) with accompanying *P*-values, derived from a general linear model.

| a) | $\beta$ (SE) | $T$-value | P-value |
| --- | --- | --- | --- |
| Density intercept | 0.046 (0.003) | 15.040 | <0.001 |
| Hourly mating probability | -0.001 (0.003) | -1.796 | 0.088 |
| Linear plasticity | >0.001 (0.004) | 0.049 | 0.961 |
| Non-linear plasticity | -0.004 (0.004) | -1.159 | 0.260 |

## Discussion

Here, we studied within-life density-dependence of male mating success in a wild insect population monitored using automated RFID-surveillance technology. We detected individual differences in average hourly mating success (i.e. intercept of the reaction norm) and density-dependent plasticity in mating success (i.e. slope of the reaction norm), and found that survival costs and fitness benefits depend on the reaction norm describing how hourly mating success changed with local density. Specifically, males that are successful in attracting mating partners under low density (of competitor males) lived longer and have higher total mating success.

### Population-level density-dependent changes in mating success

Our results show the presence of an Allee effect (Stephens et al., 1999) in mating success at the population level: a male’s hourly mating success increases, up to a certain point, when local male density increases. This type of “mate finding Allee effect” has been studied before, though studies reported mixed results (Fauvergue, 2013; Gascoigne, Berec, Gregory, & Courchamp, 2009). The most likely mechanism for the presence of an Allee effect is, in our case, that the larger acoustic choruses of males might attract proportionally more females (Gerhardt & Huber, 2002). Indeed, the relative number of females is shown to increase with increasing local density in chorusing species of frogs (Ryan et al., 1981, but see Gerhardt & Huber 2002). Our results also show that the hourly mating success starts to decline at high densities. This pattern might be caused by increasing male-male acoustic competition; under high local densities females are not able to discriminate between males due to masking interference of acoustic signals, potentially leading to suboptimal mating decisions (Gerhardt & Huber, 2002). Nevertheless, with our data we cannot separate the actual mechanisms underpinning the observed patterns in population level hourly mating success. Indeed, detailed information about the mechanisms, e.g. density-dependent acoustic competition or masking interference between males, is the next step to further our understanding about factors contributing to density-dependent mating success in the wild.

### Individual differences in mating success

Male crickets differ in the local density where their hourly mating success is highest, implying individual differences in optimal local density. Although the existence of different mating strategies have been studied before in crickets (Cade, 1981; Pascoal et al., 2014; Simmons, Tinghitella, & Zuk, 2010), mating success has not been clearly linked to variation in dynamic local competitive environments before. The traits underpinning these strategies are mostly likely related to acoustic signaling, which is the key information channel for crickets about their social environments (Gerhardt & Huber, 2002). Thus, the next step in our understanding of the evolutionary ecology of density-dependent plasticity in mating success is to study individual level investment in acoustic signaling across dynamic local densities while adopting a reaction norm approach. Interestingly, individual differences in optimal local density should lead males to select densities that best match with their mating success (Edelaar et al., 2008). However, our results show that males do not seem to select, on average, a local density in which their expected mating success is highest. There might thus be constraints preventing crickets from actively choosing the optimal social environment. Crickets most likely cannot monitor the structure of the social environment very far efficiently (Gerhardt & Huber, 2002), in which case it is hard for the focal male to determine whether the social environment is “better” somewhere else. Given that the local social density is, furthermore, very dynamic and changeable, the temporal costs of monitoring the structure of the local density might override its benefits. Finally, burrows act as shelters against predation in crickets and thus leaving the burrow is costly (Gawałek, Dudek, Ekner-Grzyb, Kwieciński, & Sliwowska, 2014). This cost might override, at least partly, the potential benefits of actively seeking the optimal local density.

### Costs and benefits related to individual level mating success

Sexual signals increasing the ability to attract opposite sex partners have been shown to be costly in many ways (Godin & McDonough, 2003; Zuk & Kolluru, 1998), potentially negatively affecting longevity. However, our results indicate that males who have high average hourly mating success do not live shorter in the wild. This pattern is somewhat consistent with the results where the expression of traits related to reproduction (e.g. song activity, mate search, dominance) do not come with clear longevity costs in the wild in *Gryllus campestris* (Rodríguez-Muñoz et al., 2019). Longevity costs related to mating success seem to be more complex and depend on the degree (and form) of density-dependent plasticity in hourly mating success. Indeed, males that have high mating success under high densities live less long than males that are more successful under low densities, contrasting our initial prediction. In other words, males that are highly competitive live less long compared to potentially less competitive males. When faced with high local densities, males most likely have to invest proportionally more in acoustic signaling in order to have high mating success, leading to higher predation and parasitism risk (Zuk & Kolluru, 1998). Indeed, in a related field cricket species, *Gryllus integer*, parasitoid wasps orient towards signaling males leading to higher parasitism rates compared to non-signaling males (Cade, 1975). Moreover, there is also evidence that “high quality” cricket males, which might have high mating success in high densities, die young because they invest heavily in sexual display (Hunt et al., 2004).

Our results also indicate that the fitness benefits, i.e. total amount of time spent with females, are higher in males that are generally successful in attracting mating partners (i.e. high average hourly probability to be with a female; intercept), but also in males who are successful in low density environments (i.e. negative linear slopes). Thus, selection overall acted on two components of the reaction norm independently, exemplifying the importance of applying the reaction norm approach in evolutionary studies of labile traits (see below). Indeed, our study is among the first ones to study selection acting on specific components of the reaction norms in the wild (Nicolaus, Brommer, Ubels, Tinbergen, & Dingemanse, 2013). Because average hourly mating success did not differ between males that do well in high versus low density environments, the total mating success is higher in males that are successful in low density because they live longer. Thus, the survival costs are lower and the fitness benefits higher for males expressing specific type of linear density-dependent plasticity in hourly mating success (i.e. negative linear slopes).

### Conclusion

Here, we show that both the intercept and slope of the reaction norm describing the mating success as a function of the local density of competitors were under selection independently: males with high average hourly mating success, but also males with high hourly mating success in low density, were selected for. Notably, we were only able to reveal the complex biology underpinning density-dependent mating success since we applied the reaction norm framework; studying selection acting on plastic traits, which vary across environmental gradients, requires specific statistical frameworks. Applying such a framework is important as it allows empiricists to quantify and study selection acting on the mean trait expression (i.e. reaction norm intercept), but also its plasticity (i.e. reaction norm slope), simultaneously and independently. Very few studies quantify selection on independent reaction norm components (but see Nicolaus et al., 2013) probably because the collection of the data required to estimate such reaction norms is extremely demanding, e.g. large amount of repeated measurements on mating success, for each individual, across naturally occurring environmental gradient is required (*sensu* Dingemanse et al., 2010; Nussey et al., 2007). In the future, we encourage empiricists to focus on studying evolutionary ecology of plastic traits by using reaction norm approach since it allows answering more diverse biological questions, potentially with less bias.

## Data Availability Statement

The data will be available in DRYAD after acceptance of the manuscript.

## Acknowledgements

P.T.N. was supported by the DFG (Deutsche Forschungsgemeinschaft; NI 1539/2-1), S.T. and N.J.D. were supported by the Ludwig-Maximilians-Universität München. Authors have no conflict of interest.

## Authors’ contributions

P.T.N. and N.J.D. contributed to the design of the study, drafting and revising of the article. P.T.N. collected and analyzed all the data. P.T.N. and S.T. contributed to the extraction of the data to a format where it can be analyzed. All authors approved the final version of the manuscript.

---

**Supplementary Table 1.** Sources of variation in hourly mating success for different local densities (i.e. 100-500 centimeter radius from a focal burrow) with distance from the edge included in the models; we present fixed (β) and random (σ^2^) parameters with 95% Credible Intervals, as well as model AIC-values, derived from univariate mixed effect models. Estimates and 95% Credible Intervals are derived by simulating the model 1000 times.

|  | Density 100 | Density 200 | Density 300 | Density 400 | Density 500 |
| --- | --- | --- | --- | --- | --- |
| **Fixed effects\*** | $\beta$ (95% CI) | $\beta$ (95% CI) | $\beta$ (95% CI) | $\beta$ (95% CI) | $\beta$ (95% CI) |
| Intercept | -4.04 (-5.94, -2.06) | -3.59 (-5.41, -1.79) | -2.97 (-4.65, -1.16) | -2.83 (-4.73, -0.99) | -2.75 (-4.59, -0.84) |
| Age | 2.32 (1.24, 3.28) | 2.52 (1.54, 3.48) | 2.34 (1.23, 3.40) | 2.10 (1.12, 3.02) | 2.21 (1.20, 3.21) |
| Age<sup>2</sup> | <-0.01 (-0.30, 0.26) | <-0.01 (-0.28, 0.28) | -0.04 (-0.29, 0.25) | -0.06 (-0.32, 0.20) | -0.04 (-0.33, 0.23) |
| Distance | -0.55 (-1.66, 0.54) | -0.52 (-1.74, 0.61) | -0.51 (-1.76, 0.68) | -0.75 (-1.97, 0.51) | -0.68 (-1.84, 0.53) |
| Density_among | -3.29 (-6.29, -0.41) | 2.41 (-0.05, 4.70) | 2.08 (0.30, 3.84) | -0.22 (-2.11, 1.67) | 0.05 (-1.76, 2.09) |
| Density_among<sup>2</sup> | 4.92 (1.60, 8.09) | 2.55 (-1.69, 6.87) | -0.67 (-3.30, 1.92) | -2.25 (-5.17, 0.71) | -1.42 (-3.64, 0.75) |
| Density_within | 0.08 (-0.08, 0.25) | 0.03 (-0.10, 0.16) | -0.33 (-0.48, -0.19) | -0.36 (-0.52, -0.22) | -0.05 (-0.24, 0.14) |
| Density_within<sup>2</sup> | -0.13 (-0.19, -0.07) | -0.12 (-0.20, -0.04) | -0.25 (-0.33, -0.16) | 0.06 (-0.03, 0.14) | -0.25 (-0.34, -0.15) |
| **Random Effects** |  |  |  |  |  |
| *Variance* | $\sigma^2$ (9% CI) | $\sigma^2$ (SD) | $\sigma^2$ (SD) | $\sigma^2$ (SD) | $\sigma^2$ (SD) |
| Residual | 3.29 (fixed) | 3.29 (fixed) | 3.29 (fixed) | 3.29 (fixed) | 3.29 (fixed) |
| Burrow | 15.48 (11.96, 19.96) | 15.37 (11.82, 19.80) | 15.49 (12.01, 19.72) | 15.62 (12.19, 19.92) | 14.81 (11.82, 18.64) |
| Date | 9.05 (6.91, 12.31) | 9.28 (7.15, 12.69) | 9.36 (7.29, 12.95) | 8.86 (6.90, 12.02) | 8.94 (6.88, 12.55) |
| Individual | 1.89 (1.43, 2.74) | 1.48 (1.17, 2.02) | 1.86 (1.46, 2.51) | 2.67 (2.07, 3.71) | 2.84 (2.22, 3.93) |
| *Model fit* | AIC | AIC | AIC | AIC | AIC |
|  | 4447.4 | 4459.2 | **4405.6** | 4456.9 | 4446.3 |

